# Non-clinical evidence supports anti-inflammatories as more effective medication than antihistamines against tarantula local effects envenomation

**DOI:** 10.1101/2021.04.13.439720

**Authors:** Bruno Ricardo Alves, Rafael Sutti, Pedro Ismael da Silva, Rogerio Bertani, Jair Guilherme Santos-Junior, Thomaz Augusto Alves Rocha e Silva, Alessandra Linardi

## Abstract

**Background:** Tarantulas are the most common invertebrates pets, especially in North America and Europe. The most commercialized genera are from Southern Asia and tropical Americas, represented by *Vitalius*, found in Southeastern Brazil, and *Brachypelma*, common in Mexican desert. Bites by these spiders in humans occurs during manipulation and generally result in clinical manifestations such local pain, erythema and oedema, with the possibility of secondary local infection. Hence, the cases are usually treated with prescription free drugs such as antihistamines and anti-inflammatories.

**Methods:** In this work, we investigated the post treatment with commercial nonsteroidal and steroidal anti-inflammatories and anti-histamines administered by oral and intraperitoneal routes on rat paw oedema induced by venoms of *V. dubius* and *B. smithi*. Hydroplethysmometer standard oedema measurement and Evans blue extravasation were performed. Dose standardization experiments showed that the *V. dubius* is more potent than *B. smithi*, and doses were established at 30 μg/paw and 60 μg/paw respectively.

**Results:** The oral post-administration of ketoprofen (non-selective cyclooxygenase inhibitor) and prednisolone (steroidal anti-inflammatory) markedly reduced a paw oedema evoked by only *V. dubius* venom, but loratadine (H_1_-antihistamine) had negligible effect on rat paw oedema induced by both venoms. Intraperitoneal administration, ketoprofen (20 mg Kg^−1^) and loratadine (5 mg Kg^−1^) reduced the rat paw oedema induced by *V. dubius* and *B. smithi* while methylprednisolone (10 mg Kg^−1^) only inhibited the oedema induced by *V. dubius*.

**Conclusion:** These results suggest that the pos-treatment with nonsteroidal and steroidal anti-inflammatory drugs are more potent than antihistamines in attenuating the local effect induced by *V. dubius* and *B. smithi* venoms.

## Background

Spiders of the family Theraphosidae (suborder Mygalomorphae) are better known in many parts of the world as tarantulas and are worldwide commercialized as pets [1]. The tropical *Vitalius* and *Brachypelma* are one of the most common genus found with breeders in North America and Europe. *Vitalius dubius* occurs in the southern part of the Brazilian state of Minas Gerais and in the state of São Paulo [2]. *Brachypelma smithi*, one of the most popular tarantulas of the world also named mexican redknee tarantula, is found along the central Pacific coast of Mexico, from southern coastal Jalisco to north-western Oaxaca State and inland to the states of Mexico and Morelos [3].

The most common defensive behavior of species of such genera is the release of abdominal urticating hairs that elicit mild allergic reactions and seldom evolve to serious injuries [4]. Thereby, this friendly profile encourages breeders to manipulate the animals and bites occasionally occurs. Despite their large size and potentially large venom yields, bites by Theraphosidae spiders in humans generally result in only local clinical manifestations such local pain, erythema and oedema, but do not induce local necrosis or systemic effects [5, 6, 7, 8]. Besides the risk of infection, inflammation and swealling at the puncture site can arise. Based on the nine patients, Isbister [6] described that the main clinical effect of bites by Australian theraphosid spiders is severe local pain, usually with puncture marks, but general systemic effects such as nausea and vomiting are uncommon. The puncture wounds from the spider’s fangs require local wound care, monitoring for signs of infection, short-term analgesia and sometimes tetanus vaccine [9]. Such cases rarely lead to medical assistance, what makes it an underestimated casuistic. Nevertheless, spider collectors often assume to have enough knowledge to realize that symptomatic healing is enough for treatment and appeal to over-the-counter medication. More, many animals are obtained through illegal market, what leads breeders to avoid notifications of such accidents.

In this work, we propose to investigate the effect of oral and intraperitoneal pos-treatment of clinical used nonsteroidal and steroidal anti-inflammatories and antihistamines on the rat paw oedema and plasma extravasation induced by venoms of *Vitalius dubius* and *Brachypelma smithi*.

## Methods

### Animals

Male Wistar-Hanover rats (200-250 g) were obtained from Department of Physiological Sciences at Santa Casa de São Paulo School of Nursing and Medicine (Faculdade de Ciências Médicas da Santa Casa de São Paulo, FCMSCSP) and were housed 5/cage at 23°C on a 12 h light/dark cycle, with free access to food and water. The experimental protocols were approved by an institutional Committee for Ethics in Animal Experimentation (CEUA/FCMSCSP, protocol n°. 001/13) and the general ethical guidelines for animal use established by the Brazilian Society of Laboratory Animal Science (SBCAL, formerly the Brazilian College for Animal Experimentation - COBEA) and EU Directive 2010/63/EU for animal experiments were followed.

The *Vitalius dubius* and *Brachypelma smithi* spiders were acquired from field collection and the Instituto Butantan, respectively. This work was performed following the CITES recommendations and national licenses required (IBAMA SISBIO 58786-1 and CGen 010115/2014-5).

### Venom and drugs

Venoms were obtained according to Rocha e Silva [10] for both species. Evans blue dye was purchased from Sigma Chemical (St. Louis, MO, U.S.A.). Ketoprofen was purchased from Sanofi-Aventis Farmacêutica (Susano, SP, Brasil). Prednisolone was purchased from Mantecorp Indústria Química e Farmacêutica (Rio de Janeiro, SP, Brazil). Methylprednisolone was obtained from União Química Farmacêutica (São Paulo, SP, Brazil). Loratadine was purchased from Medley Indústria Farmacêutica (Campinas, SP, Brazil). Formamide was obtained from Dinâmica Química Contemporânea (Diadema, SP, Brazil) and isoflurane was from Cristália (Itapira, SP, Brazil).

### Paw oedema assay

The rats were anaesthetised with isoflurane (inhaled). A subplantar injection of *V. dubius* venom (10 or 30 μg/paw) and *B. smithi* (30 or 60 μg/paw) were made into one paw on the rat. The paw volume was assessed immediately before venom, for the basal measurement and 30, 60, 120, 150 and 180 min after administration of venom, using a plethysmometer (model 7150, Ugo Basile). The final volume injected in the paw was always 0.1 ml. The same volume of saline was used in a control group. The increase in paw volume (ml) was calculated as the difference between the basal and final volumes [11]. N = 6 was used for all groups.

### Anti-inflammatory and antihistaminic activity

After standardization of dosing, groups were divided in two experimental lines based on the route of application: intraperitoneal and oral routes. Four groups of rats were envenomed and received saline (control group, 1 mL Kg^−1^), ketoprofen (20 mg Kg^−1^), methylprednisolone (steroidal anti-inflammatory; 10 mg Kg^−1^) or loratadine (5 mg Kg^−1^) by intraperitoneal route. Another four groups were also envenomed and orally treated with saline (control group, 1 mL Kg^−1^), ketoprofen (non-selective cyclooxygenase inhibitor; 50 mg Kg^−1^), prednisolone (steroidal anti-inflammatory; 20 mg Kg^−1^) or loratadine (H1-antihistaminic drug; 10 mg Kg^−1^). The equivalent corticosteroids prednisolone and methylprednisolone were used as the adequate preparation for the pharmacokinetics of the route of administration. In both experimental lines all drugs and saline were administered 15 min after respective venom inoculation.

### Vascular permeability assay

The plasma extravasation measured by Evans blue assay was performed in a restricted group of animals because this assay implies in animal sacrifice and animal welfare committee recommended as maximum of thirty animals. Hence, the most prominent results obtained with paw oedema were transposed to this assay, following the same protocol of drugs administration. Evans blue (25 mg/kg, 2.5% w/v in 0.45% NaC1) was injected intravenously immediately before subplantar injection of *V. dubius* venom (30 μg/paw). Control groups received saline by the same route as the drugs. Two hours later, the animals were killed and dye exudate were measured [12, 13]. Briefly, animal paws were amputed at the tarsocrural joint and weighed on an analytical balance. Subsequently, each paw was chopped into small pieces and placed in a test tube with formamide 4 ml and then incubated in a water bath 57^0^C for 24 h. The absorbance of the supernatants was measured at 619 nm and the concentration of Evans blue present in the extracts was determined from a standard curve of the dye prepared in formamide. The amount of Evans blue dye was expressed as μg/g of paw and the results presented as the difference in the quantity of Evans blue between the oedematogenic paw and contralateral paw. N = 5 for venom control and 4 for each treatment.

### Statistical analysis

Results are presented as means ± SD. The results were analyzed by the one way ANOVA for repeated measure considering as factor the period of oedema measured: 30, 60, 120, 150 and 180 min or one way ANOVA considering as factor the treatment. Fisher *post-hoc* was used when necessary. Values with p<0.05 were considered statistically significant.

## Results

### Paw oedema induced by V. dubius and B. smithi venoms

Intraplantar injection of *V. dubius* venom (10 and 30 μg/paw) in the rat hind-paw caused a dose and time-dependent oedema (Fig. 1), when compared to Saline groups. The one way ANOVA for repeated measures shows that there were differences in the group [F(2,13)=71.66], in the time factors [F(5,65)=108.99], as well as in the group *vs*. time interaction [F(10,65)=20.15]. For further studies, *V. dubius* venom was administered at the dose of 30 μg/paw. We also observed a dose and time-dependent oedema induced by *B. smithi* venom (30 and 60 μg/paw; Fig 1), when compared to Saline group. The ANOVA for repeated measures detected significant differences in the group [F(2,14)=69.77], the time factors [F(5,70)=103.93] and in the group *vs*. time interaction [F(10,70)=25.84]. *B. smithi* venom was administered at the dose of 60 μg/paw for subsequent studies.

**Figure 1.**
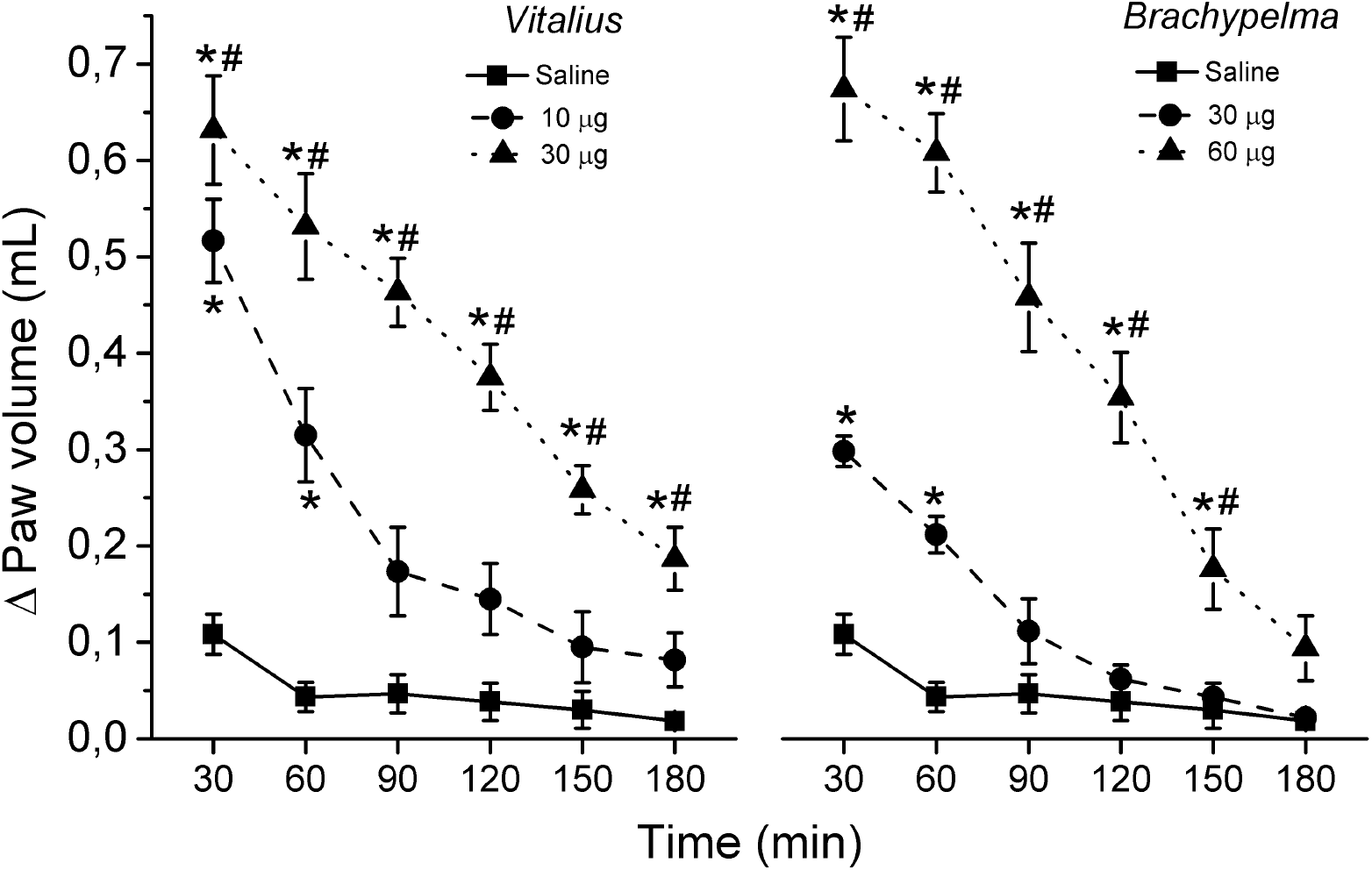
Dose dependent hind paw oedema evoked by *V. dubius* and *B. smithi* venoms. Note the higher potency of the *Vitalius* venom. *p<0,05 compared to saline; #p<0,05 compared to other dose.

Table 1 summarizes the statistical outcomes of post-treatment experiments, presented in the following items.

**Table 1.**
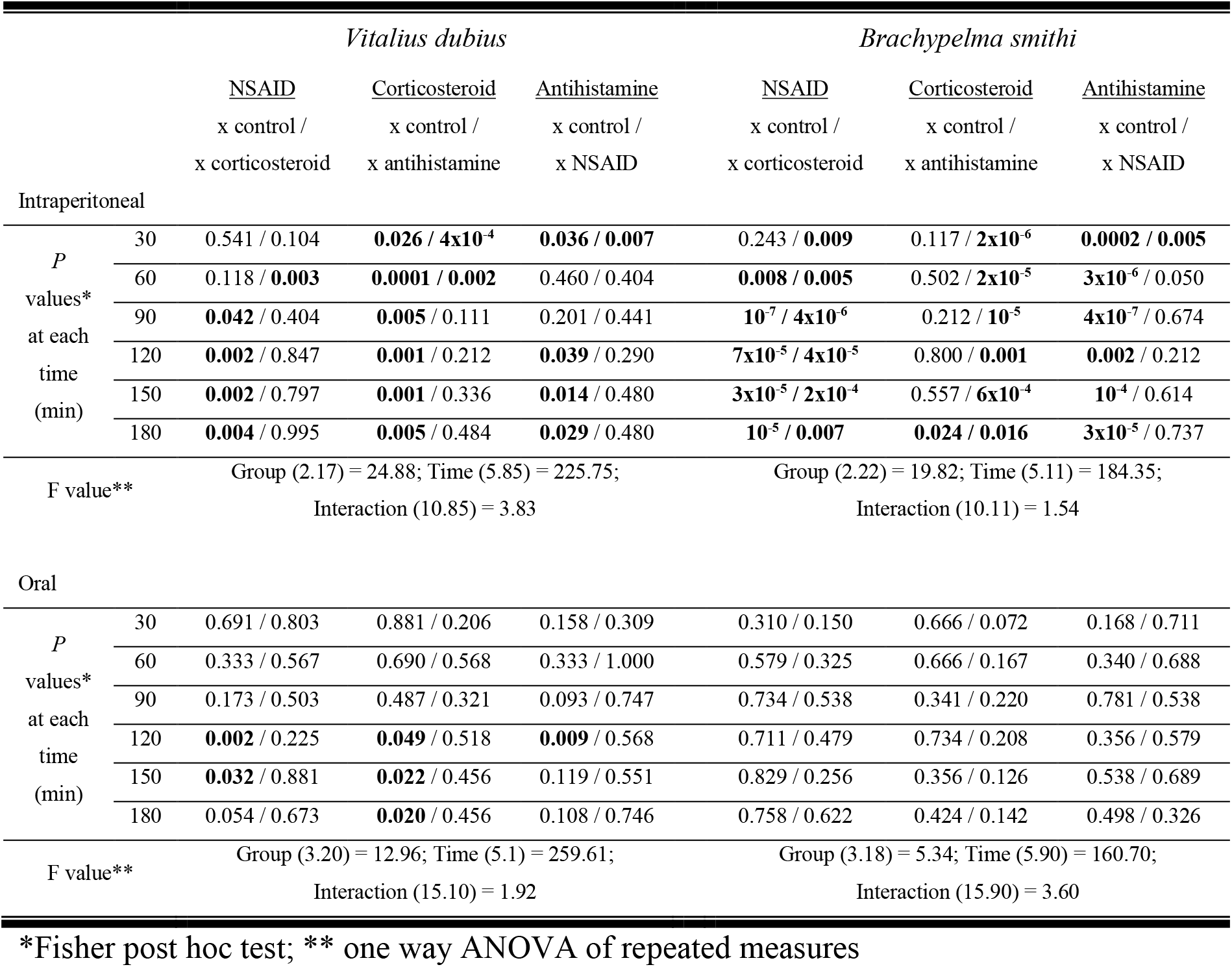
Summary table of statistics outcomes comparing different venoms, treatments, routes and times for the hind paw oedema. Columns shows the *P* values of the comparison of underlined treatment with the other two, respectively separated by bars. The significances of *P* values are in bold and indicated in figures 2 and 3.

### Effect of intraperitoneal administration (i.p.) of ketoprofen, loratadine and prednisolone on paw oedema

Figure 2 illustrates the effect of intraperitoneal post-treatment (15 min after venoms injection) of ketoprofen (20 mg Kg^−1^), loratadine (5 mg Kg^−1^) and methylprednisolone (10 mg Kg^−1^) on paw oedema induced by *V. dubius* venom (Fig. 3A) and *B. smithi* venom (Fig. 3B). For *V. dubius* venom, the repeated measures ANOVA shows that there were differences in the group and time factors and the group *vs*. time interaction (Table 1). Regarding *B. smithi* venom, same statistical analysis also describes differences in the group and time factors but in the group *vs*. time interaction there was no significant difference (Table 1). The Fisher *post-hoc* test shows that the intraperitoneal pos-administration of methylprednisolone significantly reduced the paw oedema induced by *V. dubius* venom at all subsequent intervals from 30 min, when compared to Control group. In addition, ketoprofen and loratadine significantly reduced the paw oedema in 90 and 120 min towards the end of experiment, respectively. Otherwise, significantly reduction in oedema evoked by *B. smithi* venom was not observed with methylprednisolone, but loratadine from 30 and ketoprofen from 60 until 180 min.

**Figure 2.**
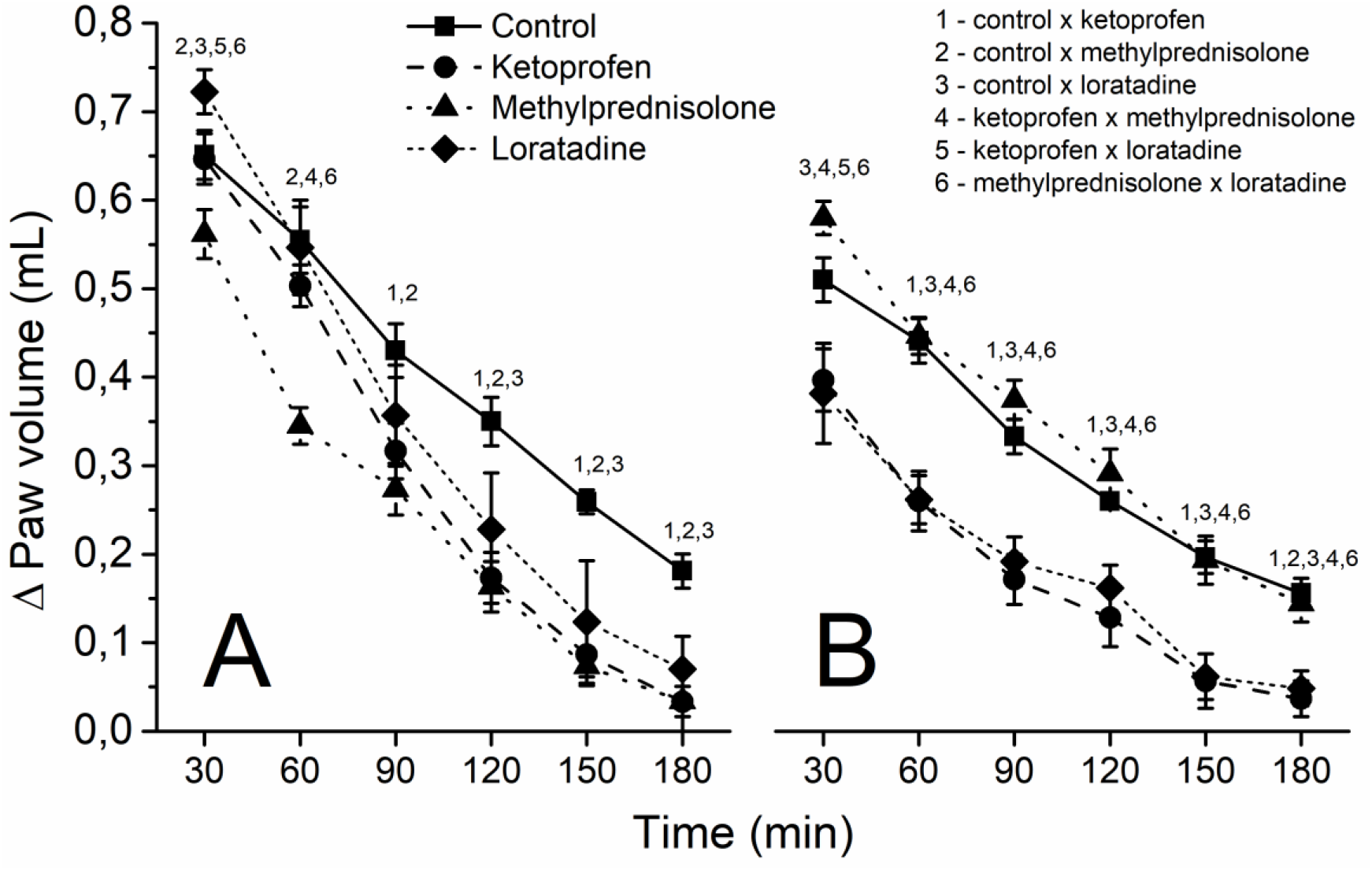
Hind paw oedema evoked by *V. dubius* (A, 30 μg) and *B. smithi* (B, 60 μg) venoms in animals treated 15 minutes after envenomation with intraperitoneal saline (control group, 1 mL Kg^−1^), ketoprofen (10 mg/kg), methylprednisolone (20 mg/kg) or loratadine (5 mg/kg). Legends are valid for both A and B graphics. Number indicates p<0,05 of respective comparison.

**Figure 3.**
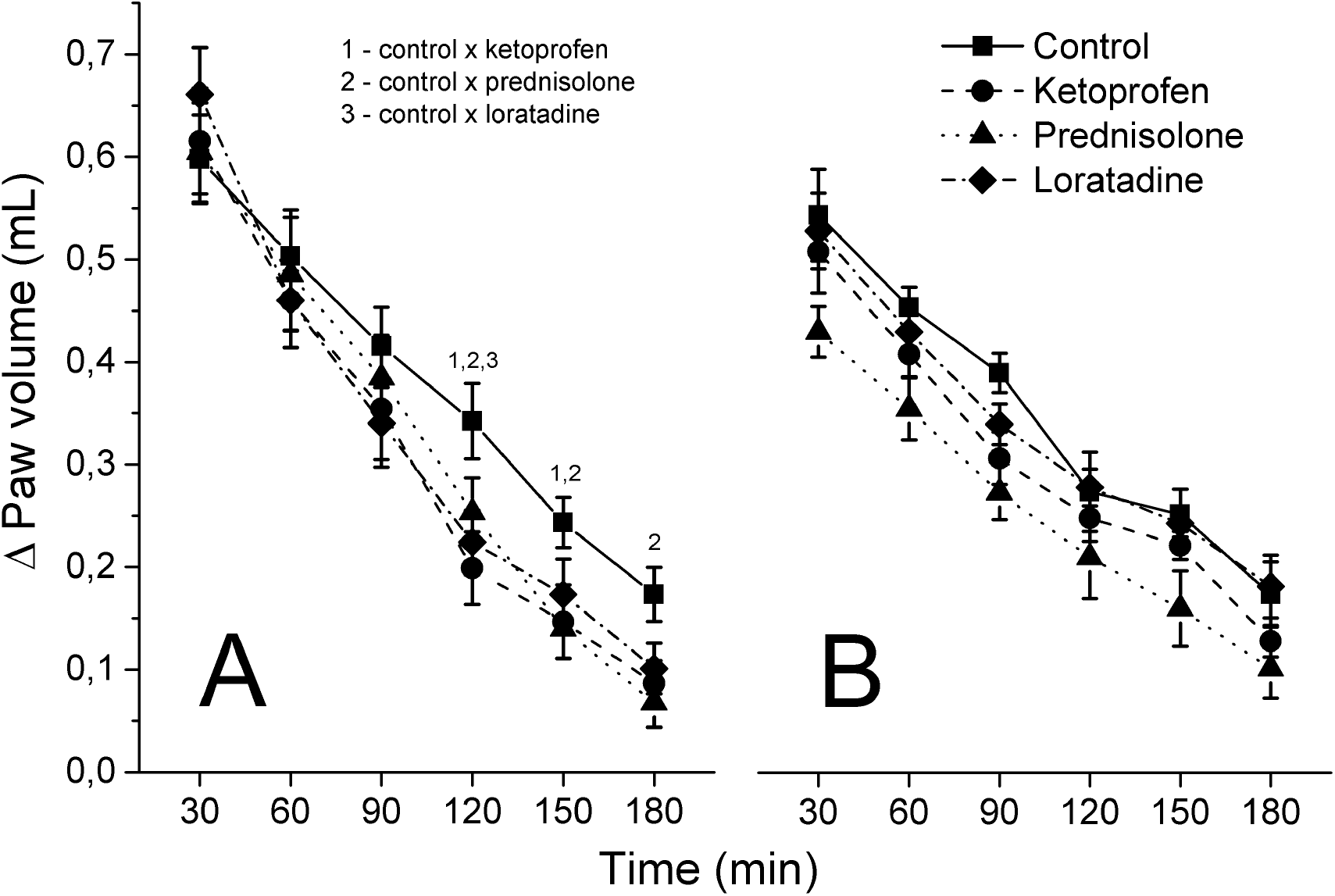
Hind paw oedema evoked by *V. dubius* (A, 30 μg) and *B. smithi* (B, 60 μg) venoms in animals treated 15 minutes after envenomation with oral saline (control group, 1 mL Kg^−1^), ketoprofen (50 mg/kg), prednisolone (20 mg/kg) or loratadine (10 mg/kg). Legends are valid for both A and B graphics. Number indicates p<0,05 of respective comparison.

### Effect of oral administration (p.o.) of ketoprofen, loratadine and prednisolone on paw oedema

Figure 3 shows the effect of oral pos-treatment (15 min after venoms into the paw) of ketoprofen (50 mg Kg^−1^), loratadine (10 mg Kg^−1^) and prednisolone (20 mg Kg^−1^) on paw oedema induced by *V. dubius* venom (Fig. 4A) and *B. smithi* venom (Fig. 4B). For *V. dubius* venom, the repeated measures ANOVA shows that there were differences in the group and time factors, as well as in the group *vs*. time interaction. However, for *B. smithi* venom, there were differences only in the group and time factors, but there were no significant differences in the group *vs*. time interaction (Table 1). This indicates that oral prednisolone and ketoprofen are able to diminish the oedema caused by *B. smithi* venom as a hole treatment and an increase in the number of animals may reach significance in group *vs*. time interaction. The Fisher *post-hoc* test shows that the oral post-treatment with ketoprofen significantly reduced the paw oedema induced by *V. dubius* venom at intervals of 120 and 150, when compared to Control group. However, the prednisolone post-treatment significantly reduced the paw oedema only at intervals from 120 min towards the end of experiment. Loratadine significantly alter the paw oedema induced by *V. dubius* venom only at 120 min.

**Figure 4.**
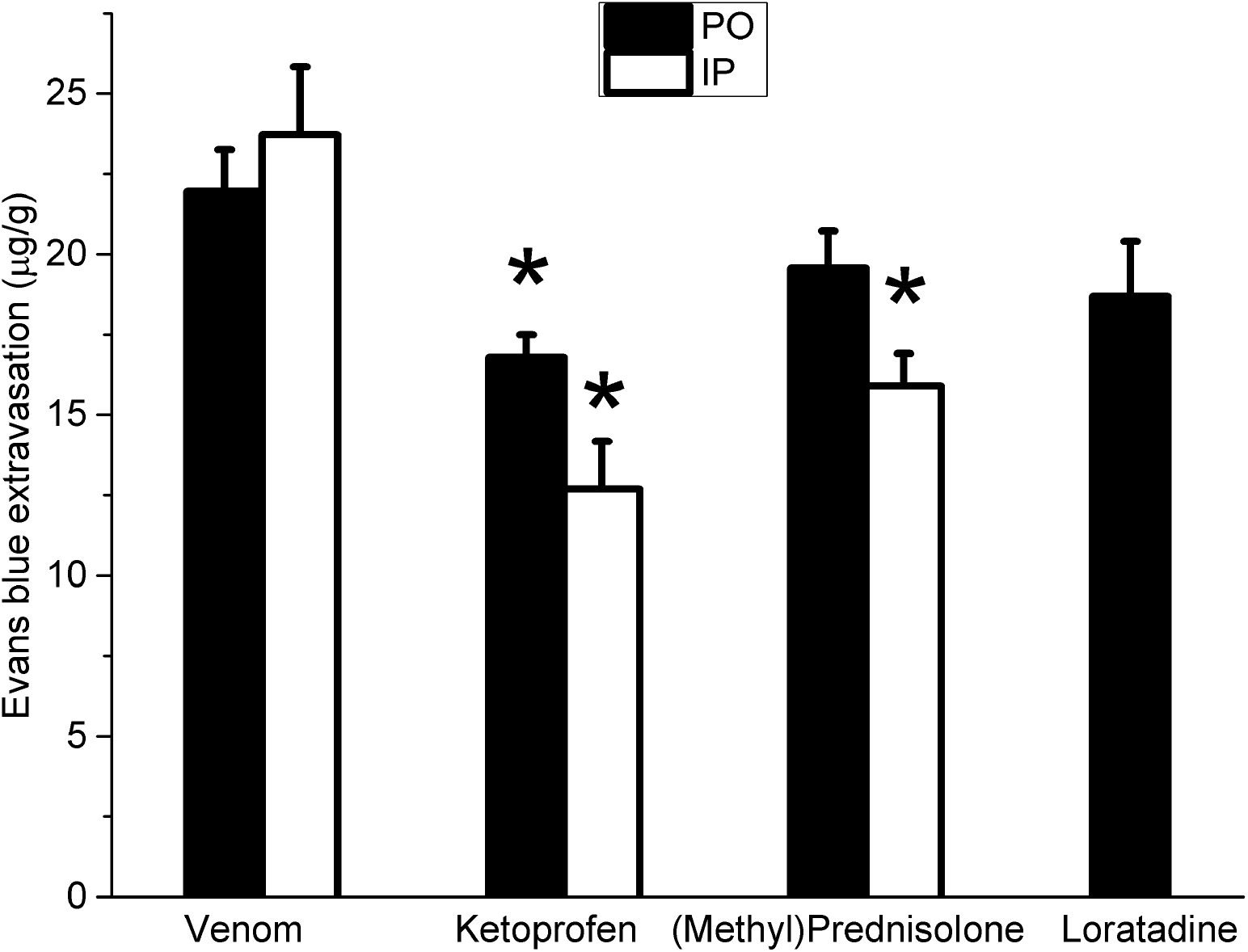
Effect of oral (p.o., black bars) or intraperitoneal (i.p., white bars) administration of saline (venom group), ketoprofen (20 mg/kg i.p and 50mg/kg; p.o), methylprednisolone (10 mg/kg; i.p.)/prednisolone (20 mg/kg; p.o.) and loratadine (5 mg/kg; i.p.) on Evans blue extravasation induced by *V. dubius* venom after 120 minutes of venom injection. Each column represent mean ± standard error (N = 5 for venom control and 4 for each treatment). *p<0,05 *vs*. control (venom alone)

### Effect of intraperitoneal (i.p.) or oral (p.o.) administration of ketoprofen, loratadine and prednisolone or methylprednisolone on vascular exudation induced by V. dubius venom

The time interval of 120 min was chosen to evaluate the Evans blue extravasation induced by *V. dubius* venom in rats. The one way ANOVA detected significant differences in the groups [F(2.14)=10.8] after intraperitoneal post-treatment, as well as it was observed differences between groups [F(2,11)=12.10] after oral post-treatment (Fig. 4). The Fisher *pos-hoc* test showed that the intraperitoneal treatment of ketoprofen (20 mg Kg^−1^) and methylprednisolone (10 mg Kg^−1^) significantly reduced the Evans blue extravasation induced by *V. dubius* venom, when compared to Control group. Regarding the oral post-treatment, Fisher *pos-hoc* test detected that the ketoprofen (50 mg Kg^−1^) and prednisolone (20 mg Kg^−1^), but not loratadine (5 mg Kg^−1^), significantly inhibited the Evans blue extravasation, when compared to Control group.

## Discussion

American tarantula spiders are usually not aggressive and easily manipulable often living in regions with easy access for being captured. Despite the concern of conservationists about the increase of illegal commerce, these spiders have been kept as pets mainly in North America and Europe, but also in many other parts of the world. Nevertheless, stressed or hungry animals may display defensive behavior such as the release of urticating hairs or even biting an unsuspected owner.

The bite of the tarantulas is quite painful mainly due to the size of the fangs, but few cases of envenomation are reported, probably due to low toxicity to humans [5, 8] and therefore it is underreported to health systems. Hence, mild clinical consequences such as swelling and moderate pain rarely requires medical assistance, mainly amongst experienced breeders that usually appeal to over the counter medicines.

Although reports of minor effects in humans, a number of studies have demonstrated significant toxicity of theraphosid venoms in various animals, including rats, mice, cats, birds and dogs [14, 15, 16]. In addition, it is also recognized that theraphosid spiders can cause more severe effects in domestic animals, including death [6, 17, 18].

The venom composition of the two species considered in this work has been studied and share characteristics such as the presence of hyaluronidase and lack of phospholipase and proteinolytic activity [10, 19]. *V. dubius* is the only specie of the genus that has already published data on venom composition, including an ionotropic blocker polyamine with yet antimicrobial activity [20, 21]. The described oedema activity [22] was not associated to a specific toxin since it was demonstrated to be induced by various pathways. Otherwise, the venoms of the *Brachypelma* genus have been widely studied, comprising insecticidal and neurotoxic activity [23, 24], with small molecules related to tissue toxicity [25]. Despite scarce macromolecules, theraphosid spiders produce a wide range of peptides that can differ between species of the same genera and usually have selective ligand properties [26] that may also elicit inflammation and local reactions.

Our results demonstrated that the venom of *V. dubius* and *B. smithi* induced a paw oedema in rats, with difference in the respective doses, what evidences a higher potency of the South American spider. The local effect of the venom of the *V. dubius* was first inhibited by the corticosteroid, followed by ketoprofen and loratadine, respectively, when drugs were administered intraperitoneally. Concerning the oral route, the efficacy of loratadine was statistically lower than other drugs when compared to control group. These observations meet previous results concerning the pharmacology of the oedema caused by *V. dubius* that showed no participation of histamine and bradykinin, but a significant action of serotonin, eicosanoids, neurokinins and nitric oxide [22].

On the other hand, the local action of the *B. smithi* venom have suffered intense inhibition by loratadine and ketoprofen but not by methylprednisolone when administered intraperitoneally. The oral route for the same drugs, although observed only as a group effect, conversely demonstrated oedema inhibition by corticosteroid, followed by ketoprofen and lack of loratadine effect, opening the room for the discussion about pharmacological differences of the venoms. The results indicate a possible role of histamine in the oedema caused by *B. smithi* venom, although this component was not found in the composition of previously studied venom [25]. Comparable paw oedema was observed with *B. epicureanum* but no pathways or toxins were investigated [27]. These results are evidences of the presence of peptides that may elicit inflammation in this venom.

The plasma extravasation measured by Evans blue assay was performed only with *Vitalius* venom because of the higher potency compared to *Brachypelma*. Loratadine was tested orally to check if its efficacy was indeed lower, since the intraperitoneal administration demonstrated oedema healing. Surprisingly, ketoprofen was the most efficient drug on avoiding plasma extravasation, with significative reduction through both routes of administration, while corticosteroid was significant only through intraperitoneally administration. Finally, oral loratadine or corticosteroid were not different from control, evidencing a primary role of cyclooxygenase derivates in plasma extravasation evoked by *V. dubius* venom.

There are few studies related to the mechanisms involved in local manifestations in accidents caused by theraphosid spiders. It was demonstrated that tarantulas may produce pro-inflammatory and nociceptive peptides acting as capsaicin receptor agonist and pain-related sodium channel activators, respectively [28, 29]. Our group demonstrated the participation of neurokinins, cyclooxygenase and nitric oxide pathways on skin oedema caused by *V. dubius* venom [22]. It is remarkable that most of the studies that investigate the mechanisms of action of venom and/or its fractions, administered drugs before the envenomation, i.e. before the onset of the inflammatory response by venom [30, 31, 32]. Otherwise, the importance of post-treatment studies of envenomation was recently corroborated. The systemic effects of various *Poecilotheria* genera venoms has been studied in mice and demonstrated several neurological and muscular symptoms. The authors demonstrated that post-treatments with at least three drugs were able to avoid cramps and movement stereotytpy [33].

Also, a comparison between intraperitoneal and oral routes arises. It was observed that intraperitoneal administration of drugs was more effective when compared the oral administration, especially when considering the paw oedema induced by *B. smithi* venom. Oral administration is considered to be safe, efficient and easily accessible with minimal discomfort to the patient compared to other routes of administration. However, the absorption of the drug may be reduced due to digestive enzymes, gastrointestinal tract motility, pH, and especially by the hepatic first-pass metabolism. It is known that antihistamines are extensively transformed in liver to inactive metabolites or a minor fraction of active ones such as desloratadine [34]. Otherwise, the pharmacokinetic pathways of (methyl)prednisolone are complex and protein bound distribution and interconversion between prednisolone and prednisone [35] may explain the lack of inhibition of *B. smithi* venom when administered intraperitoneally.

The use of NSAIDs such as ketoprofen are often related to pain control, but our work demonstrated the benefits of these drugs in avoiding the local effect, as a second hand effect. Also, systemic (i.e. oral) corticosteroids showed a significative action against envenomation. However, concerning the side effects of NSAIDs and corticosteroids, the first is usually related to stomach discomfort in a short term use [36], while the second shall be avoided through systemic routes because of potential effects on immune system [37], since local infection is a concern during tarantula envenomation. The action of loratadine may be useful in cases which is the only available drug or with the arousal of a systemic allergic reaction, but the liver first pass metabolism may not allow a locally effective concentration.

## Conclusion

The main contribution of the present work is the comparison of three commercially available drugs to treat local oedema caused by tarantulas envenomation. Our results showed that anti-inflammatory medicines are more effective in healing local effects than antihistamines. If the systemic (oral) strategy is chosen, NSAIDs may be a better choice, corticosteroids shall be used topically and antihistamines reserved for the case of allergic reactions.

## Abbreviations

ANOVA: analysis of variance
*B. smithi*: *Brachypelma smithi*
CEUA: Committee for Ethics in Animal Experimentation
CGen: Commission for Biodiversity Genetic Access
COBEA: Brazilian College for Animal Experimentation
COX: cyclooxygenase
FCMSCSP: Santa Casa de São Paulo School of Medicine
H1: histamine receptor 1
i.p.: intraperitoneal
IBAMA: Brazilian Institute for Environment
NSAIDs: non steroidal anti-inflamatory drugs
p.o.: oral administration
SBCAL: Brazilian Society of Laboratory Animal Science
SD: standard deviation
SISBIO: Biological Sampling License System
*V. dubius*: *Vitalius dubius*

## Aknowledgements

The authors would like to thank Fundação Florestal de São Paulo, the managers of PETAR, Intervales and Serra do Mar-Curucutu State Parks and Centro de Controle de Zoonoses de Itu.

## Availability of data and materials

The datasets generated during and/or analyzed during the current study are available from the corresponding author on reasonable request.

## Authorship Contributions

*Participated in research design*: AL, JGSJr, TAARS and BRA

*Conducted experiments*: BRA, RS and AL

*Contributed new reagents or analytic tools*: RB, PISJr, TAARS and RS

*Performed data analysis*: JGSJr, AL and TAARS

*Wrote or contributed to the writing of the manuscript*: AL, TAARS, JGSJr, PISJr and RB

## Ethics approval

The experimental protocols were approved by an institutional Committee for Ethics in Animal Experimentation (CEUA/FCMSCSP, protocol n°. 001/13) and the general ethical guidelines for animal use established by the Brazilian Society of Laboratory Animal Science (SBCAL, formerly the Brazilian College for Animal Experimentation - COBEA) and EU Directive 2010/63/EU for animal experiments were followed.

